# Endosomal trafficking defects alter neural progenitor proliferation and cause microcephaly

**DOI:** 10.1101/2020.08.17.254037

**Authors:** Jacopo A. Carpentieri, Amandine Di Cicco, David Andreau, Laurence Del Maestro, Fatima El Marjou, Laure Coquand, Jean-Baptiste Brault, Nadia Bahi-Buisson, Alexandre D. Baffet

## Abstract

Primary microcephaly and megalencephaly are severe brain malformations defined by reduced and increased brain size, respectively. Whether these two pathologies arise from related alterations at the molecular level is unclear. Microcephaly has been largely associated with centrosomal defects, leading to cell death. Here, we investigated the consequences of *WDR81* loss of function, which cause severe microcephaly in patients. We show that WDR81 regulates endosomal trafficking of EGFR, and that loss of function leads to reduced MAP kinase pathway activation. Mouse radial glial progenitor cells knocked-out for *WDR81* display reduced proliferation rates, leading to reduced brain size. These proliferation defects are rescued *in vivo* by the expression of megalencephaly-causing mutated Cyclin D2. Our results identify the endosomal machinery as an important regulator of RG cell proliferation rates and brain growth. They demonstrate that microcephaly and megalencephaly can be due to opposite effects on the proliferation rate of radial glial progenitors.

## Introduction

Development of the neocortex relies on neural stem cells called radial glial (RG) cells, that generate the majority of cortical neurons^1^. Neuronal production is restricted to a short period during which all excitatory neurons are produced^2^. This leaves little room for compensatory mechanisms to occur, and alterations during this critical period lead to brain malformations^3^. Indeed, the developing neocortex is highly sensitive to perturbations, and a large number of mutations have been described to specifically alter its growth, but not that of other organs^4^.

Primary microcephaly is a severe neurodevelopmental disorder characterized by a head circumference that is more than 3 standard deviations (SD) below the mean^5^. The major molecular cause of microcephaly lies in defects in centrosomal number^6,7^, maturation^8^ and mitotic spindle regulation^9-11^, leading to apoptotic cell death. In fact, apoptosis appears to be the leading cause of microcephaly in animal models, irrespective of the upstream affected molecular pathway^12-14^. Reduced proliferation rates of progenitors, while proposed to be a putative cause of microcephaly^15^, has received much less experimental support. One notable example is the gene encoding IGFR1, which is mutated in syndromic forms of microcephaly, and when deleted in mouse leads to reduced proliferation and small brain size^16,17^.

On the opposite end of the spectrum, megalencephaly (MEG) is a neuronal disorder characterized by brain overgrowth (3 SD over the mean)^18^. The causes of megalencephaly are diverse, but activating mutations in the Pi3K-AKT-mTOR and the Ras-MAPK pathways have been identified as important underlying events^19-21^. Mouse and cerebral organoid models for these activated pathways demonstrated increased proliferation of radial glial cells leading to tissue overgrowth^22-24^. Stabilizing mutations in the downstream target Cyclin D2 were also reported, and its ectopic expression in mouse brain stimulated progenitor proliferation^25^.

The EGF receptor (EGFR) and its ligands are major regulators of tissue growth^26^. Accordingly, knock-out of EGFR leads to a dramatic atrophy of the cerebral cortex^27^. Progenitor cells appear to become responsive to EGF at mid-neurogenesis, while at earlier stages they rather exhibit FGR2 dependence^28^. Endosomal trafficking of EGFR plays a major role in the regulation of its activity: while most EGFR signaling is believed to occur at the plasma membrane, internalization of EGFR is critical for signal termination^29^. Following endocytosis, internalized cargos follow different trafficking routes including recycling towards the plasma membrane or delivery to lysosomes for degradation^30^. Phosphatidylinositols (PtdIns) are major regulators of this process, defining endosomal compartment identity. Early endosomes are characterized by the presence of the small GTPase RAB5 and PtdIns3P, and late endosomes by RAB7 and PtdIns(3,5)P_2_^31^. Recently, WDR81 and its partner WDR91 were shown to act as negative regulators of class III phosphatidylinositol 3-kinase (PI3K)-dependent PtdIns3P generation, therefore promoting early to late endosomal conversion^32^. In WDR81 knock-out (KO) HeLa cells, endosomal maturation defects led to delayed EGFR degradation^32^.

We recently reported compound heterozygous mutations in the human *WDR81* gene, that result in extreme microcephaly associated with reduced gyrification of the neocortex^33^. Here, we generated a mouse knock-out model that largely recapitulates the human phenotype. Mutant brains are not only smaller but also display altered neuronal layering. We demonstrate that microcephaly is the result of reduced proliferation rates of radial glial progenitors, but not of cell death. Mechanistically, we show that WDR81 mutation delays EGFR endosomal trafficking and leads to reduced activation of the MAPK signaling pathway. These proliferation defects can be rescued by expressing a megalencephaly-causing mutated cyclin D2, indicating that microcephaly and megalencephaly can be due to opposite effects of the proliferation rates of radial glial cells.

## Results

### WDR81 KO mice display reduced brain size and altered neuronal positioning

In mice, two WDR81 isoforms have been identified. A long isoform (210 kDa) encompassing an N-terminal BEACH domain, a central transmembrane region, and a C-terminal WD40 repeat domain; and a shorter isoform (81 kDa) lacking the BEACH domain (**Figure 1A**). Measurements of mRNA isolated from embryonic E14.5 cortex extracts indicated that the long WDR81 isoform was highly dominant, with only trace levels of the short isoform (**Figure 1B**). We generated two WDR81 KO mice using gRNAs targeting the beginning of exon 1 (KO-1, affecting isoform 1), and the end of exon 1 (KO-2, affecting both isoforms) (**Figure 1A)**. Both lines displayed frameshifts leading to the appearance of a premature STOP codon (**Figure S1A**). QPCR measurements in KO1 did not reveal any upregulation of isoform 2, indicating an absence of compensation (**Figure 1B**). Moreover, a strong reduction of isoform 1 mRNA levels was observed, likely due to non-sense mRNA decay (**Figure 1B**). WDR81 homozygote mutant embryos and pups were detected at sub-mendelian rates, and did not live for more than 21 days (**Figure S1B**).

**Figure 1.**
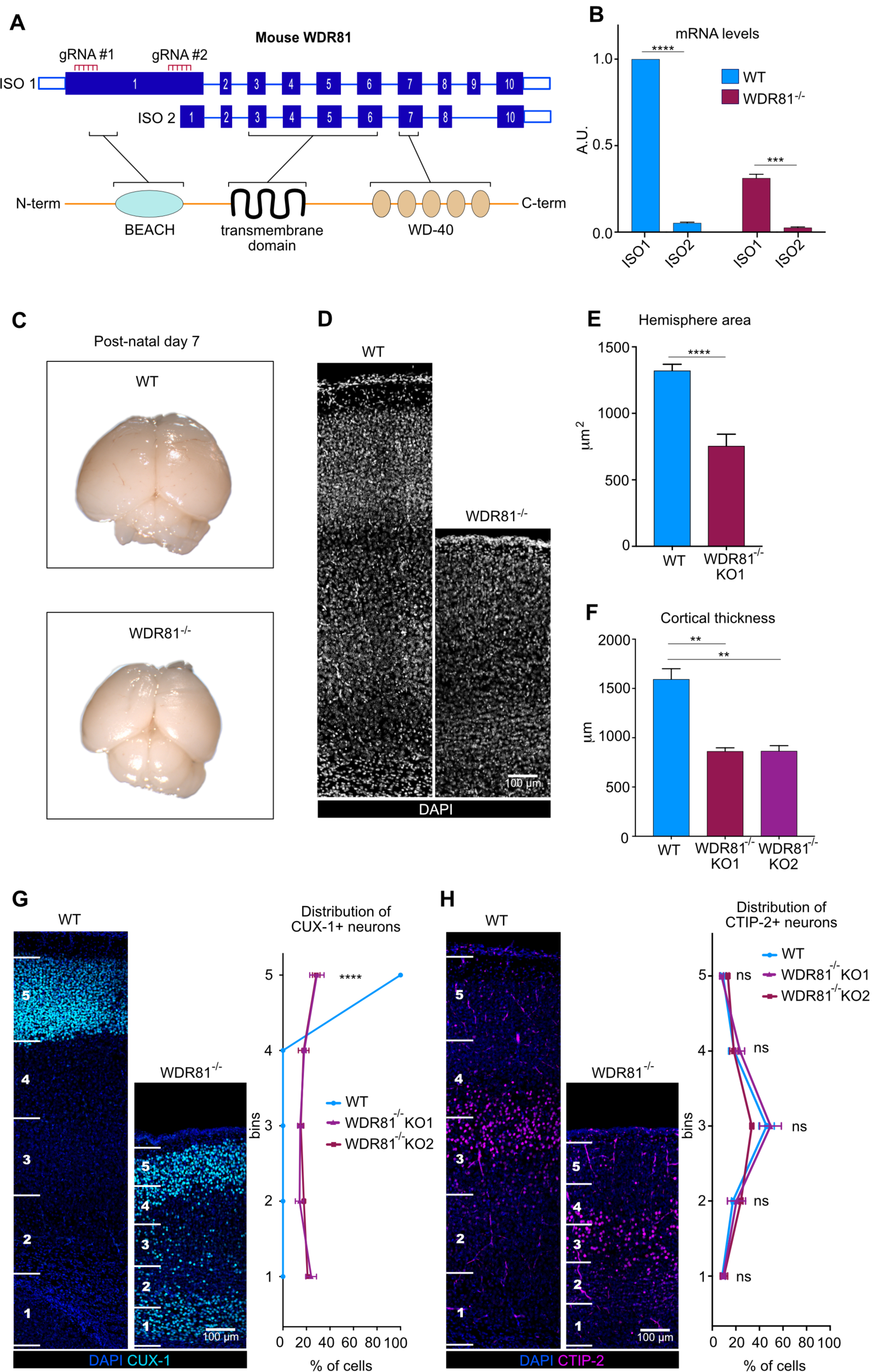
WDR81 KO mice display reduced brain size and altered neuronal positioning. **A.** Schematic representation of mouse WDR81 isoforms and predicted structure. **B**. Quantification of WDR81 isoforms 1 and 2 mRNA levels in WT and WDR81^-/-^ E14.5 cortices (n = 3 brains per genotype). Isoform 1 (ISO1) is the dominant isoform and its levels are strongly reduced in WDR81^-/-^ cortices. **C**. WDR81^-/-^ postnatal day 7 brains are microcephalic and display reduced cortical surface area as compared to WT brains. **D**. DAPI staining of P7 WT and WDR81^-/-^ cross sections reveals reduced cortical thickness in mutants. **E**. Quantification of hemisphere areas in P7 WT and KO1 brains (n = 4 brains per genotype). **F**. Quantification of cortical thickness in P7 WT, KO1 and KO2 brains (n = 3 brains per genotype). **G**. CUX-1 staining in P7 WT and WDR81^-/-^ cortices. Quantification of CUX1+ neuronal positioning reveals dispersion throughout the thickness of the neocortex (n = 3 brains per genotype). **H**. CTIP-2 staining in P7 WT and WDR81^-/-^ cortices. Quantification does not neuronal positioning defects, with CTIP-2+ neurons still concentrated in the third bin (n = 3 brains per genotype). (**G, H)** WT and mutant cortices were divided into 5 bins of equal size to measure neuronal relative positioning, independently of cortical thickness. **p<0,01;***p<0,001; ****p<0,0001 by unpaired t-tests.

We then analyzed brain size and organization in P7 WDR81^-/-^ pups. Both KO lines were severely microcephalic, with a reduced hemisphere area (**Figures 1C and 1E**). Cortical thickness was also greatly reduced (by ∼54%), suggesting defects both in tangential and radial expansion of the brain (**Figures 1D and 1F**). We next analyzed neuronal positioning in WDR81^-/-^ P7 cortices. The localization of upper layer late-born neurons was severely affected, with a large number of CUX-1-positive neurons dispersed throughout the cortex (**Figure 1G**). Deeper neurons, which are born earlier during cortical development were however correctly positioned, as indicated by the localization of CTIP-2-positive neurons (**Figure 1H**). Overall, mice knocked out for WDR81 have reduced brain size and altered neuronal positioning, largely recapitulating the microcephaly and lissencephaly phenotypes reported in humans. These phenotypes were observed for both WDR81 KO lines and we therefore next focused our analysis on KO-1 (referred to as WDR81^-/-^ from here on).

### WDR81 KO alters radial glial progenitor proliferation

To identify the causes of reduced brain size in WDR81^-/-^ pups, we tested for alterations of neocortex development at embryonic stages. To test for proliferation defects, we first measured the mitotic index of WDR81^-/-^ radial glial progenitors, as defined by the percentage of phospho-Histone H3 (PH3)-positive cells out of total PAX6-positive cells. While at E12.5, proliferation appeared normal, mitotic index of WDR81^-/-^ radial glial progenitors was severely reduced at E14.5 and E16.5 (**Figure 2A**). Strikingly, TBR2-positive intermediate progenitors appeared to cycle normally throughout development (**Figure 2B**). Therefore, WDR81 mutation specifically alters proliferation of radial glial progenitors at mid and late neurogenic stages. We next analyzed further these proliferation defects and measured the percentage of cells in S phase. A 30-minute BrdU pulse revealed an increased amount of WDR81^-/-^ radial glial progenitors in S-phase (**Figures 2C and 2D**). To test whether this was due to a longer duration of S-phase, we performed a double BrdU-EdU pulse, in order to measure the rate of S-phase exit. Mice were first injected with BrdU, followed by a second injection with EdU 4 hours later. This assay revealed a decreased proportion of cells that exited S phase (BrdU+/EdU-) in WDR81^-/-^ brains, indicating a longer S phase in mutant radial glial progenitors (**Figures 2E and 2F**).

**Figure 2.**
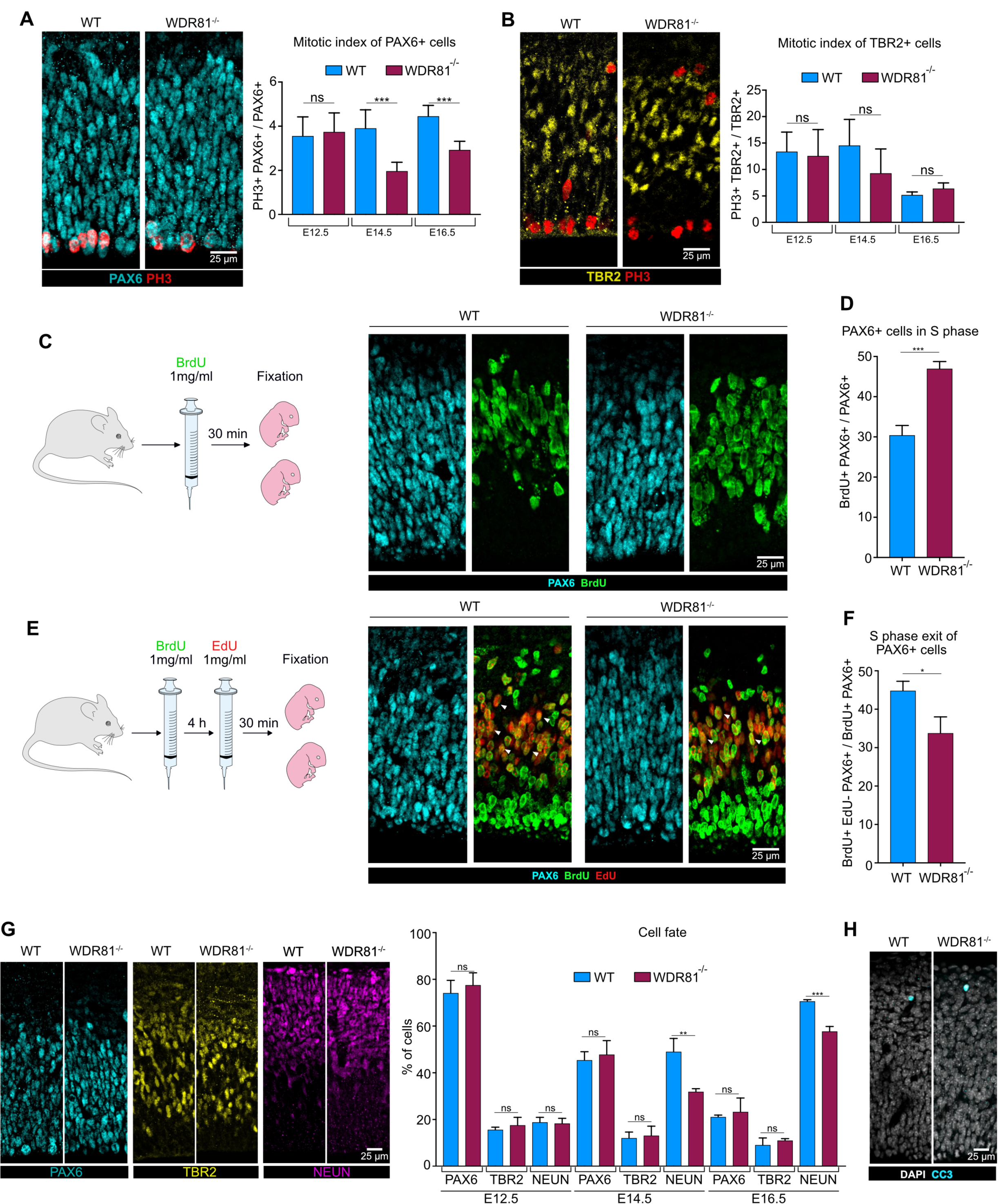
WDR81 KO alters radial glial progenitor proliferation. **A.** PAX6 and PH3 double staining in E14.5 WT and WDR81^-/-^ brains. Quantification of the mitotic index of PAX6+ cells reveals decreased proliferation of WDR81^-/-^ radial glial progenitors at E14.5 and E16.5 (n = 5-8 brains per condition). **B**. TBR2 and PH3 double staining in E14.5 WT and WDR81^-/-^ brains. Quantification of the mitotic index of TBR2+ cells indicates that proliferation of WDR81^-/-^ intermediate progenitors in not affected (n = 3 brains per condition). **C**. Schematic representation of the BrdU labeling experimental approach and Pax6 and BrdU staining in E14.5 WT and WDR81^-/-^ brains. **D**. Quantification of the percentage of BrdU+ PAX6+ out of total PAX6 cells reveals increased number of cells in S phase in WDR81^-/-^ radial glial progenitors at E14.5 (n = 3 brains per condition). **E**. Schematic representation of the BrdU / EdU double labeling experimental approach and PAX6, BrdU and EdU staining in E14.5 WT and WDR81^-/-^ brains. **F**. Quantification of the percentage of of BrdU+ EdU-PAX6+ out of the total BrdU+ PAX6+ cells reveals a decreased proportion of cells that exited S phase following BrdU injection in WDR81^-/-^ radial glial progenitors at E14.5 (n = 3 brains per condition). **G**. Staining for the cell fate markers PAX6 (radial glial progenitors), TBR2 (Intermediate progenitors) NEUN (Neurons) in E14.5 WT and WDR81^-/-^ brains, and quantification of cell fate distribution at E12.5, E14.5 and E16.5 (n = 3-5 brains per condition). **H**. Staining for Cleaved Caspase-3 (CC3) and DAPI in E14.5 WT and WDR81^-/-^ brains, showing an absence of apoptosis induction. *p<0,05; **p<0,01; ***p<0,001 by unpaired t-tests.

An alternative potential cause of reduced brain size is premature differentiation of progenitor cells. To test this, embryonic cortices were stained for PAX6 (radial glial progenitors), TBR2 (intermediate progenitors) and NEUN (neurons) at different developmental stages and the proportion of each cell population was measured. We did not observe any decrease in the proportion of progenitors out of the total cell population throughout development indicating they did not prematurely differentiate (**Figure 2G**). In fact, we even detected a reduction of the proportion of neurons at mid and late neurogenesis (**Figure 2G**). Finally, we analyzed apoptotic cell death in WDR81^-/-^ cortices. Staining for cleaved caspase-3 (CC3) did not reveal any increased apoptosis, which remained almost undetectable both in WT and mutant embryos (**Figure 2H**). Therefore, reduced brain size in WDR81^-/-^ mice is not the result of premature progenitor differentiation or increased apoptotic cell death, but appears to be a consequence of reduced radial glial progenitor proliferation rates.

### Reduced proliferation and EFGR signaling in WDR81 patient cells

We next tested whether similar proliferation defects could be observed in patient cells mutated for WDR81. Two mutant primary fibroblast lines, derived from skin biopsies, were analyzed and compared to two control fibroblast lines. The mitotic index of both patient cells was strongly decreased, mimicking the mouse radial glial progenitor phenotype (**Figures 3A and 3B**). As an alternative measurement of proliferation, cells were stained for Ki67, which also revealed a substantial decrease for both patient cell lines (**Figures 3C and 3D**).

**Figure 3.**
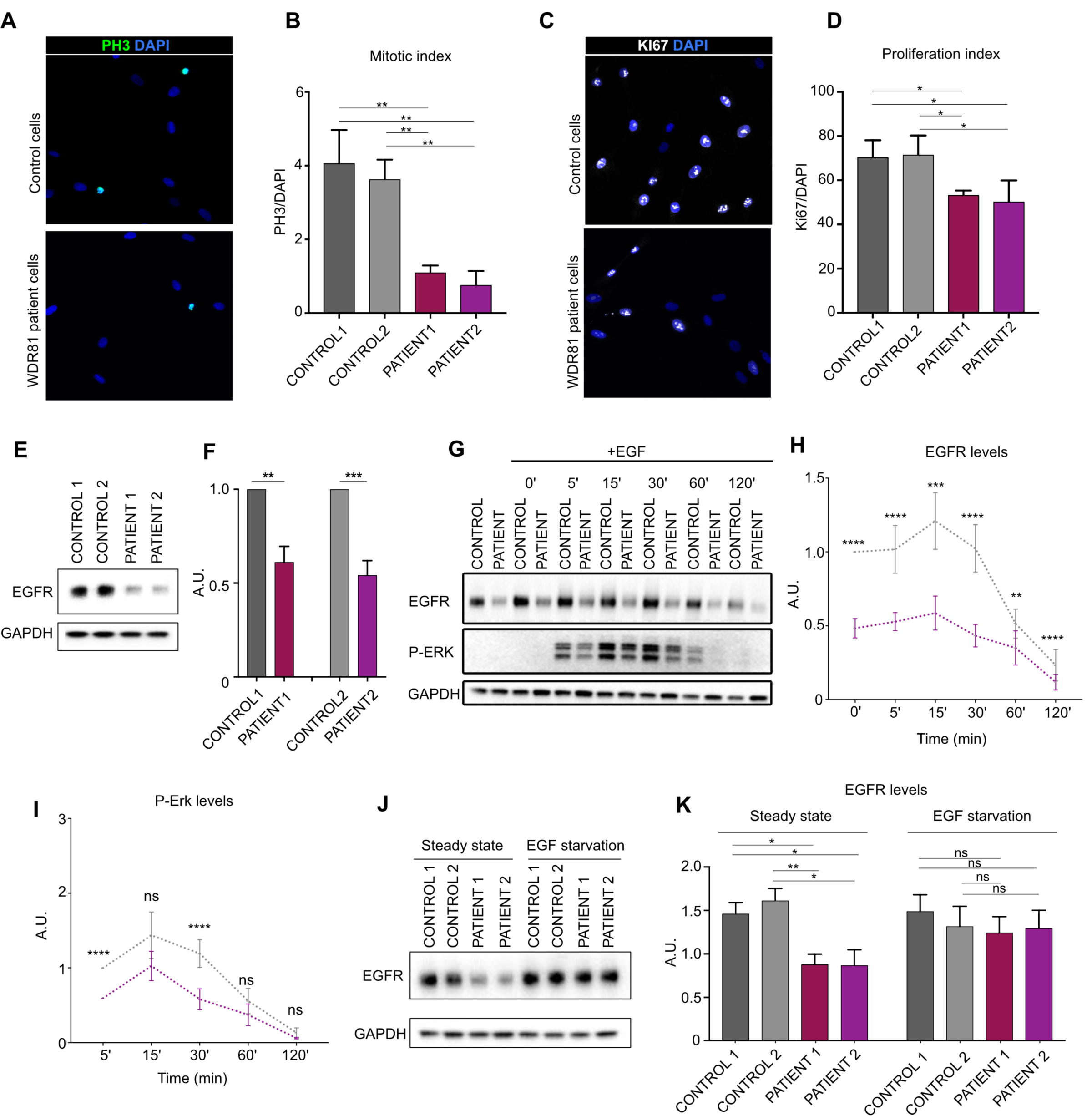
Reduced proliferation and EFGR signaling in WDR81 patient cells. **A.** PH3 and DAPI staining in control and WDR81 patient fibroblasts **B**. Quantification of the percentage of PH3+ cells reveals decreased mitotic index in patient cells (n = 3). **C**. Ki67 and DAPI staining in control and WDR81 patient fibroblasts. **D**. Quantification of the percentage of Ki67+ cells shows decreased proliferation in patient cells (n = 3). **E**. Western Blot for EGFR in control and WDR81 patient fibroblasts. **F**. Quantification reveals a strong reduction of EGFR levels in patient cells (n = 5). **G**. Time course of EGFR and P-ERK levels in control and WDR81 patient fibroblasts following an EGF pulse. **H**. Quantification of EGFR levels, normalized to control levels at T0 (n = 5). **I**. Quantification of P-ERK levels, normalized to control levels at T5 (n = 5). **J**. Western Blot for EGFR in control and WDR81 patient fibroblasts at steady state (+EGF) and cultivated for 24H in the absence of EGF. **K**. Quantification reveals a restoration of EGFR levels following starvation (n = 3).. *p<0,05; **p<0,01; ***p<0,001; ****p<0,0001 by unpaired t-tests.

In radial glial progenitors, we detected proliferation defects from mid-neurogenesis (**Figure 2A)**, which fits with the time when these cells start responding to EGF^28^. This observation suggested a potential alteration in the EGFR signaling pathway. In order to test this, we monitored the activity of this signalling pathway in control and patient fibroblasts. Strikingly, we observed that the protein levels of EGFR itself were drastically reduced in both patient cell lines (**Figures 3E and 3F**). We next measured the activation of the mitogen-activated protein kinase (MAPK) signaling pathway in response to EGF stimulation. Consistent with the decreased levels of EGFR, the phosphorylation of ERK was reduced in both patient fibroblasts following an EGF pulse (**Figures 3G-I and S2**). Therefore, WDR81 patient cells display reduced EGFR levels, leading to a reduced activation of the MAPK signalling pathway upon EGF stimulation.

The levels of EGFR are known to be tightly regulated, through complex feedback loops and the balance between recycling and degradation of the internalized receptor^34^. We therefore asked whether reduced EGFR levels were a consequence of defects within the EGFR pathway itself. To test this, we EGF-starved cells and measured the levels of EGFR 24 hours later. In patient cells, starvation rescued EGFR levels to the ones of control cells (**Figures 3J and 3K**). These results indicate that reduced EGFR levels are only seen when the pathway is activated. Given that EGFR internalization upon EGF binding is a major regulator of the pathway, this data points towards intracellular processing defects of EGFR.

### WDR81 is required for endosomal trafficking of EGFR

WDR81 is known to regulate endosomal maturation as well as autophagic clearance of aggregated proteins (aggrephagy)^32,35^. Importantly, these two functions are independent and act via the WDR81 specific binding partners WDR91 and p62, respectively. We therefore tested whether one of these factors affected neocortex development similarly to WDR81. To perform this, we *in utero* electroporated shRNA-expressing constructs for WDR81, WDR91 and p62 in E13.5 developing brains and analyzed cell distribution at E17.5. Consistent with the KO data, WDR81 knock-down (KD) strongly affected neurodevelopment (**Figures 4A and 4B**). In particular, a large fraction of KD cells accumulated in the intermediate zone (IZ), at the expense of the germinal zones and cortical plate. This phenotype was phenocopied by WDR91 KD, but not by p62 KD which did not appear to affect cell distribution (**Figures 4A and 4B**). These results support the endosomal function of WDR81 as a critical player for proper neocortex development.

**Figure 4.**
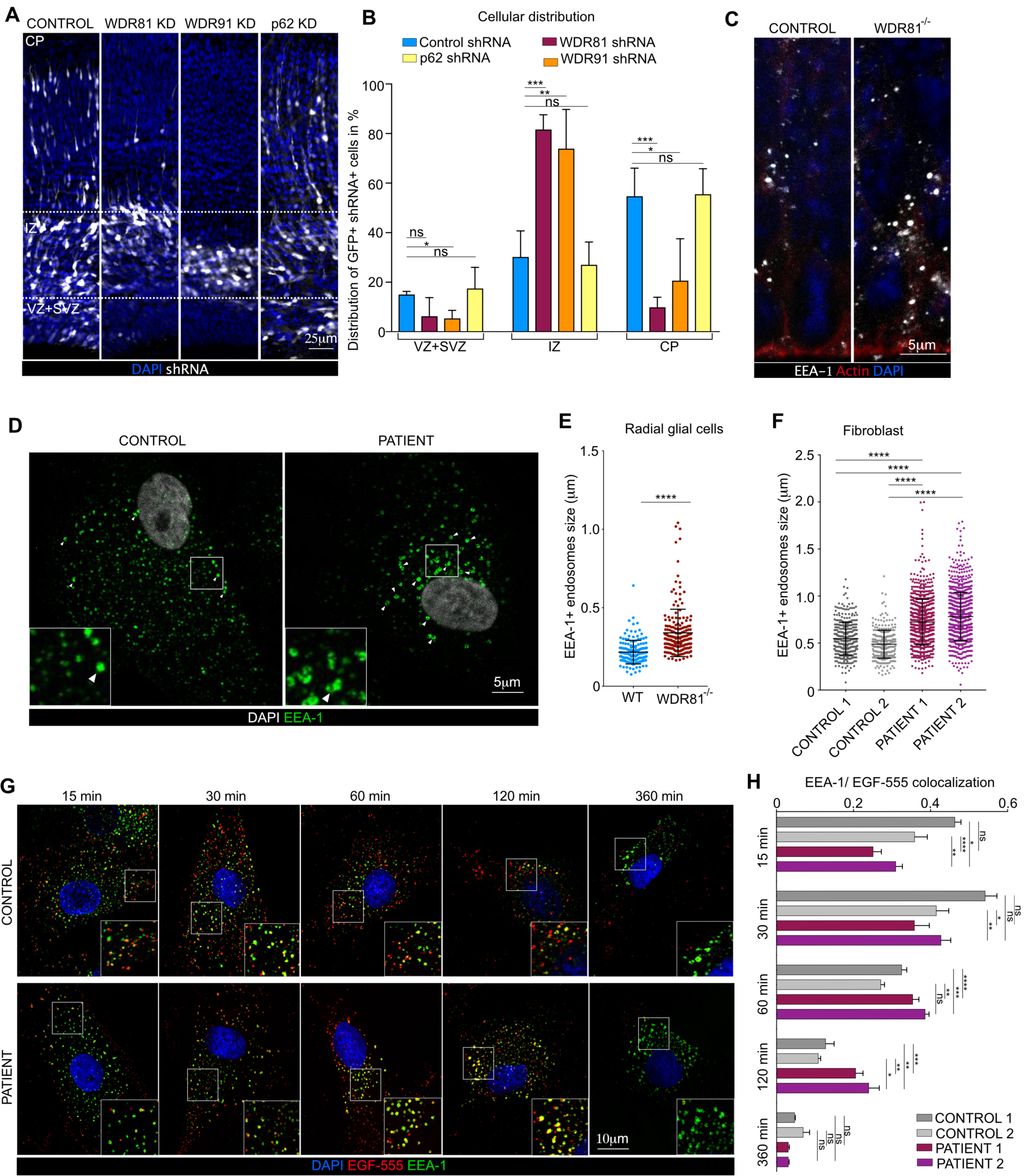
WDR81 is required for endosomal trafficking of EGFR. **A.** Expression of control shRNA and shRNA-mediated knockdown constructs for WDR81, WDR91 and p62. Plasmids were delivered by in utero electroporation at E13.5 and analysis was performed at E16.5. Ventricular Zone and Sub-Ventricular Zone (VZ +SVZ), Intermediate Zone (IZ) and Cortical Plate (CP). **B**. Quantification of electroporated cell distribution reveals major accumulation in the IZ following WDR81 and WDR91 knockdown (n = 4 brains per condition). **C**. Ventricular zone of E14.5 WT and WDR81^-/-^ mice cortices stained for EEA1 and Actin. **D**. Control and WDR81 patient fibroblasts stained for EEA1. **E**. Quantification of individual EEA1+ early endosomes in WT and WDR81^-/-^ VZ reveals increased size in mutant brains (n = 3 brains per condition). **F**. Quantification of individual EEA1+ early endosomes in control and WDR81 patient fibroblasts reveals increased size in mutant cells (n = 20 cells per condition). **G**. EGF^555^ uptake assay in control and WDR81 patient fibroblasts, and stained for EEA1. **H**. Quantification of EGF^555^ and EEA1 colocalization during EGF^555^ uptake reveals prolonged colocalization between EGF and early endosomes in WDR81 patient cells (n = 28 cells per condition). *p<0,05; **p<0,01; ***p<0,001; ****p<0,0001 by unpaired t-test (A & H) and Mann-Whitney tests (E & F).

We next tested whether endosomal defects could be observed in WDR81 KO radial glial progenitors. Staining for various endolysosomal compartments revealed a specific alteration of EEA1+ early endosomes, which appeared strongly enlarged (**Figure 4C**). Quantification of their size confirmed this observation, revealing a 63% average increase (**Figure 4E**). To test whether this is a conserved feature of WDR81 patient cells, we measured early endosome size in mutant fibroblasts. Again, EEA1+ endosomes were found to be swollen, with an increased proportion of large endosomes (>0,5 µm) (**Figures 4D and 4F**). These results are consistent with previous observation made in KO HeLa cells and demonstrating a role for WDR81 in negative regulation of Class III Pi3K^32^.

Because these endosomal defects are a potential cause of altered EGFR signaling, we next tested whether EGFR endosomal trafficking was affected in WDR81 mutant cells. Cells were first starved for 24 hours to restore EGFR to the levels of control cells, and subsequently pulsed with fluorescent EGF^555^, to monitor internalization and clearance of EGF-bound EGFR. In both patient cells, EGF^555^ was shown to accumulate longer within EEA1+ early endosomes (**Figure 4G**). Quantification of the colocalization between EGF^555^ and EEA1 revealed that this delay was particularly important 120 minutes after EGF internalization (**Figure 4H**). Therefore, WDR81 is critical for endosomal homeostasis and trafficking of internalized EGFR following EGF binding.

### Megalencephaly-causing mutation rescues progenitor proliferation in WDR81 mutant brains

Our results indicate that trafficking defects of EGFR can arise from mutations in WDR81, and lead to reduced activation of the MAPK signaling pathway. They further show that reduced radial glial progenitor proliferation is a cause of primary microcephaly. Megalencephaly is characterized by brain overgrowth and can be due to increased cell proliferation during development^18^. Major causes include gain-of-function mutations in AKT3 and its downstream target Cyclin D2^25,36^. Together, these data suggest that microcephaly and megalencephaly can be the consequence of opposite effects on the proliferation rates of radial glial progenitors. To further test this, we analyzed the effect of a megalencephaly-causing Cyclin D2^Thr280Ala^ mutant on the proliferation of WDR81 KO radial glial progenitor. Degradation-resistant Cyclin D2^Thr280Ala^, WT Cyclin D2 or a control vector were expressed using *in utero* electroporation in WT and WDR81-mutant mice brains at E 14.5, and the mitotic index of PAX6 cells was measured at E16.5. Following expression of the control empty vector, we confirmed the reduced mitotic index in WDR81^-/-^ brains (**Figures 5A and 5B**). Moreover, expression of Cyclin D2^Thr280Ala^ in WT brain increased mitotic index, indicating that this megalencephaly-causing mutation indeed stimulates radial glial progenitor proliferation rates (**Figures 5A and 5B**). Strikingly, expression of degradation-resistant Cyclin D2^Thr280Ala^ in WDR81^-/-^ brains rescued the mitotic index reduction (**Figures 5A and 5B**). WT cyclin D2 was also able to rescue proliferation, although to a lesser extent, indicating that large amounts of this protein, either due to overexpression or impaired degradation is able to restore proliferation (**Figures 5A and 5B**). Together, these results indicate that a megalencephaly-causing mutation can overcome the effect of a microcephaly-causing mutation on the proliferation of radial glial progenitors. These two pathologies can therefore arise from a highly related cause: an imbalance in cell cycle regulation leading either to reduced brain growth or to brain overgrowth (**Figure 5C**).

**Figure 5.**
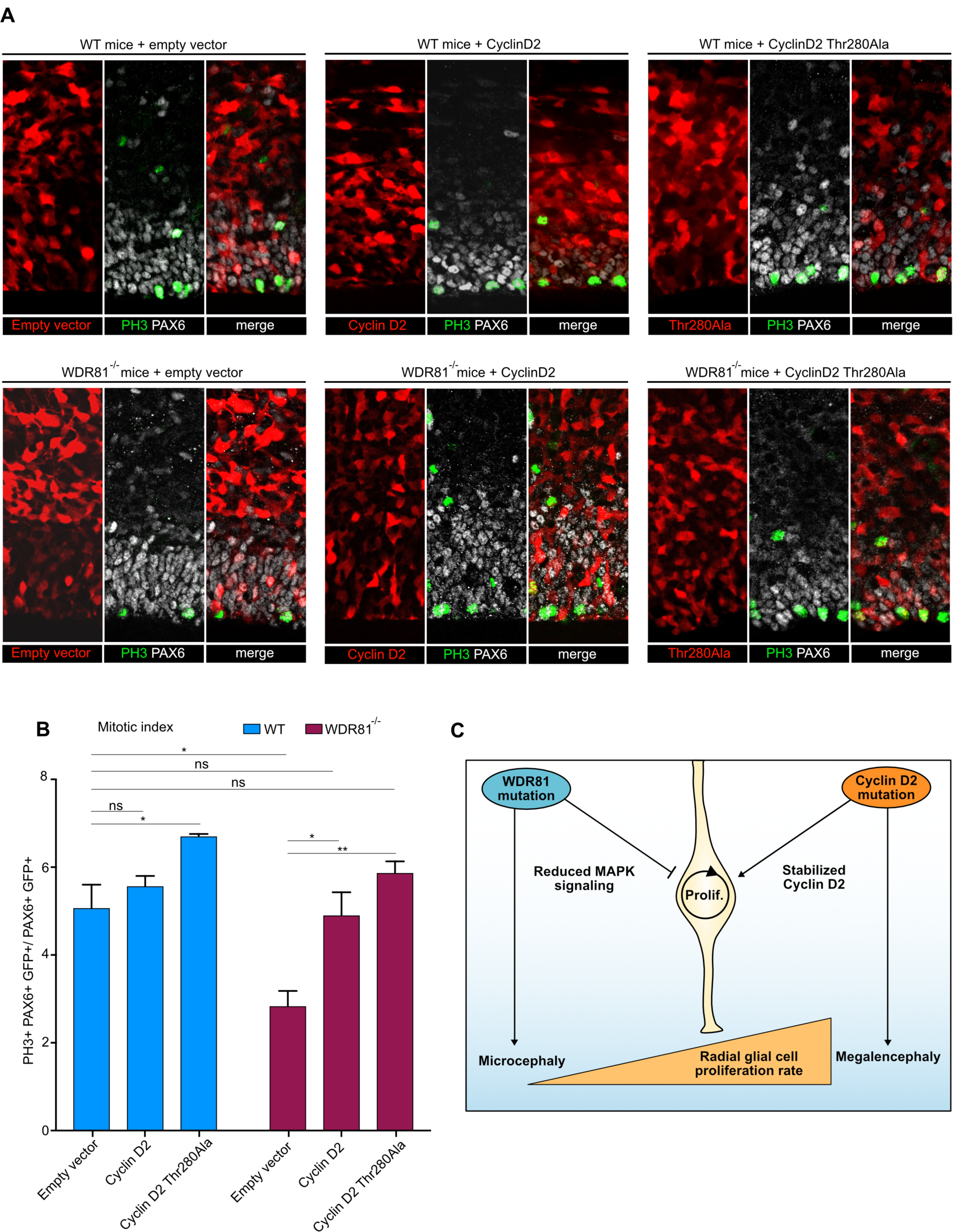
Undegradable Cyclin D2^Thr280Ala^ rescues WDR81^-/-^ proliferation index. **A.** Expression of Cyclin D2, Cyclin D2^Thr280Ala^ and empty vector in WT and WDR81^-/-^ brains. Constructs were *in utero* electroporated at E14.5, and brains were fixed at 16.5 and stained for PAX6 and PH3. **B**. Quantification of the percentage of mitotic (PH3+) electroporated radial glial cells (PAX6+) out of total electroporated radial glial cells reveals rescue of mitotic index in WDR81^-/-^ cells expressing Cyclin D2^Thr280Ala^ (n = 3 brains per condition). **C**. Model. WDR81 loss of function leads to reduced activation of the MAPK signaling pathway downstream of EGFR, to reduced radial glial progenitor proliferation, and to microcephaly. Gain of function in the Pi3K-AKT pathway or stabilizing mutations in Cyclin D2 lead to increased radial glial progenitor proliferation, and to megalencephaly. Cyclin D2 mutants can rescue proliferation defects in WDR81^-/-^ brains, indicating that these two pathologies can arise from opposite effects on the proliferation rates of radial glial progenitor. *p<0,05; **p<0,01 by unpaired t-tests.

## Discussion

In this study, we investigated the mechanisms by which mutation in the *WDR81* gene leads to severe microcephaly in patients. We show that KO mouse recapitulates many features of the phenotype previously observed in patients and that the endosomal maturation function of WDR81 is critical for neocortex development. WDR81 is required for endosomal clearance of internalized EGFR and normal activation of the mitogenic MAPK signaling pathway. In the absence of WDR81, the proliferation rate of radial glial cells is affected, leading to reduced brain size. Importantly, cell death does not appear to contribute to this phenotype. Proliferation defects can be rescued by the expression of a megalencephaly-causing mutated cyclin D2, highlighting a tight functional link between these two pathologies.

Membrane trafficking has been poorly investigated in radial glial cells, albeit its predicted implication in many important processes including cargo polarized transport, secretion of extracellular matrix components, or endocytic processing of surface receptors for lysosomal degradation or recycling. We show here that the endosomal maturation machinery plays a critical role in the processing of internalized EGFR in RG cells, and is required for their proliferation. Neurogenesis depends on EGFR activity, with radial glial cells becoming responsive to EGF from mid-neurogenesis^27,28^. Accordingly, we find that WDR81^-/-^ RG cells are specifically affected at E14,5 and E16,5 stages of development. Why the proliferation rate of IPs was not affected is unclear but EGF is secreted into the cerebrospinal fluid from the choroid plexus and apical contact may be critical for responsiveness^37,38^. EGFR was previously reported to be asymmetrically inherited during radial glial cell division, generating a daughter cell with higher proliferative potential^37^. Later in development, EGFR also acts as an important regulator of astrocyte differentiation^39^. Our data point to the intracellular processing of EGFR as an important level of control for the regulation of proliferation in radial glial cells. WDR81 is likely to affect the trafficking of other cargos^40^, which may also impact cell radial glial cell proliferation. Moreover, the trafficking of neuronal cargoes, such as adhesion molecules, is likely to lead to the altered neuronal positioning observed in KO mice, and to the lissencephaly phenotype in human.

In principle, microcephaly can be the consequence of premature progenitor differentiation, reduced proliferation rates, or cell death. While centrosomal defects leading to apoptosis have been described, reduced proliferations rates have received little experimental support^16^. In mouse, RG cells produce eight to nine neurons during a short neurogenic period, before differentiating^2^. We show here that *WDR81* mutation does not affect the modes of division of RG cells nor cell survival, but act solely through a reduction of their proliferation rate, leading to reduced brain size. This highlights the absence of compensatory mechanisms in the developing neocortex, where all neurons must be produced in a defined temporal window. During corticogenesis, G1 lengthening is associated with increased neurogenic divisions at the expense of symmetric amplifying divisions^41,42^. We did not detect such cell fate changes in WDR81^-/-^ brains. This is likely due to the fact that the proliferation rate of mutant RG cells is only affected at a stage where the vast majority of cells already perform neurogenic divisions^1^. At the macroscopic level, microcephaly and megalencephaly can appear as opposite phenotypes. However, whether they can originate from related underlying causes at the molecular level is unclear. We show here that microcephaly and megalencephaly can be due to opposite on the proliferation rates of RG cells, and can therefore be viewed as two sides of the same coin.

## Materials and Methods

### Animals

All experiments involving mice were carried out according to the recommendations of the European Community (2010/63/UE). The animals were bred and cared for in the Specific Pathogen Free (SPF) Animal Facility of Institut Curie (agreement C 75-05-18). All animal procedures were approved by the ethics committee of the Institut Curie CEEA-IC #118 and by French Ministry of Research (2016-002.

### Guide RNA selection and preparation

gRNA sequences targeting exon 1 of WDR81 have been identified and selected using the online software CRIPSOR (crispor.tefor.net). Forward and reverse oligonucleotides were annealed and cloned into px330 plasmid. To generate Cas9 mRNA and gRNA, *in vitro* transcriptions were performed on the Cas9 pCR2.1-XL plasmid and gRNA plasmids, using the mMESSAGE mMACHINE T7 ULTRA kit and the MEGAshortscript T7 kit (Life Technologies), respectively. Cas9 mRNA and sgRNAs were then purified using the MEGAclear Kit (Thermo Fisher Scientific) and eluted in RNAse-free water. The gRNA and Cas9mRNA quality were evaluated on agarose gel.

### Generation of WDR81 Knock-Out mice

Eight-week-old B6D2F1 (C57BL/6J × DBA2) females from Charles River France, were superovulated by intraperitoneal (i.p.) administration of 5 IU of Pregnant Mare Serum Gonadotropin followed by an additional i.p. injection of 5 IU Human Chorion Gonadotropin 48 h later. Females were mate to a stud male of the same genetic background. Cytoplasmic microinjection was performed into mouse fertilized oocytes using Cas9 mRNA and sgRNA at 100 ng/µl and 50 ng/µl, respectively in Brinster buffer (10 mM Tris-HCl pH 7.5; 0.25 mM EDTA). Microinjected zygotes were cultured in Cleave medium (Cook, K-RVCL-50) at 37°C under 5% CO2 and then implanted at one cell stage into infundibulum of E0.5 NMRI pseudo-pregnant females (25-30 injected zygotes per female). According to the genotyping strategy, 3 mice showed modified allele out of a total of 22 pups. The founders were then backcrossed to C57BL6/J.

### Genotyping WDR81^-/-^ animals

Mice DNA was extracted from a piece of hear (adult) or tail (dissected embryos), put at 96 degrees in lysis tampon overnight. The DNA was then amplified via PCR using WDR81 specific primers: for KO1 Forward: GGCGGAAAGTGGTTCTTACA, Reverse : AGCCACCTCCTGCATGAACC; for KO2 Forward: GGCTTGTAGTGGTTCTGTAC, Reverse : GATCCTTCTGCATTCCAA. For KO1 the amplicon was purified using the nucleospin purification kit (Machenery and Nagel) and then exposed to the restriction enzyme AfeI (New England Bioscience). The restriction enzyme only cuts the mutant DNA giving rise to two DNA pieces of 200bp. For KO2, the amplicon was sanger-sequenced using the GATC-Eurofins platform.

### Real-time reverse-transcription PCR

Wild type and WDR81^-/-^ cortices were dissected at E14.5 in 1 ml of TRIZOL (Thermo Fisher 15596026). The mRNA was isolated as follows: TRIZOL + sample solution was exposed to chlorophorm for 7 minutes at room temperature and centrifuged at 15.000g for 30 min at 4 degrees. The translucent solution formed was then transferred in 1ml of isopropanol, incubated 7 minutes at room temperature and centrifuged at 4 degrees 10.000g. The pellet of nucleic acid formed was then washed in ethanol 70% and centrifuged 5 min at 10.000g at 4 degrees. The pellet was then resuspended in water. The nucleic acids solution was purified from DNA using TURBO DNA-free Kit (Thermofisher). The mRNA obtained was then retrotranscribed using the RT reverse transcription Kit (Thermofisher). Real time RT-PCR was performed using the qPCR Master Mix kit (Thermofisher) and the WDR81 Forward/Reverse primers; GAPDH gene was used for internal control and for normalization. Primers used for WDR81 isoform 1 are forward: AGTGGATCCTTCAGACAGCC, Reverse : GAAGCCAGCCACAACACTC. Primers used for WDR81 isoform 2 are Forward: AGTGGATCCTTCAGACAGCC, Reverse : CTGACTTGTAGTGGTGCGTG

### *In utero* electroporation of mouse embryonic cortex

Pregnant mice at embryonic day 13.5 or 14.5 were anesthetized with isoflurane gas, and injected subcutaneously first with buprenorphine (0.075mg/kg) and a local analgesic, bupivacaine (2 mg/kg), at the site of the incision. Lacrinorm gel was applied to the eyes to prevent dryness/irritation during the surgery. The abdomen was shaved and disinfected with ethanol and antibiotic swabs, then opened, and the uterine horns exposed. Plasmid DNA mixtures were used at a final concentration of 1 µg/µl per plasmid, dyed with Fast Green and injected into the left lateral ventricle of several embryos. The embryos were then electroporated through the uterine walls with a NEPA21 Electroporator (Nepagene) and a platinum plated electrode (5 pulses of 50 V for 50 ms at 1 second intervals). The uterus was replaced and the abdomen sutured. The mother was allowed to recover from surgery and supplied with painkillers in drinking water post-surgery. Electroporated brain were harvested at E16.5 and E17.5.

### Immunostaining of brain slices

Mouse embryonic brains were dissected out of the skull, fixed in 4% Pfa for 2 hours, and 80 µm-thick slices were prepared with a Leica VT1200S vibratome in PBS. Slices were boiled in citrate sodium buffer (10mM, pH6) for 20 minutes and cooled down at room temperature (antigen retrieval). Slices were then blocked in PBS-Triton X100 0.3%-Donkey serum 2% at room temperature for 2 hours, incubated with primary antibody overnight at 4°C in blocking solution, washed in PBS-Tween 0.05%, and incubated with secondary antibody overnight at 4°C in blocking solution before final wash and mounting in aquapolymount. Image analysis, modifications of brightness and contrast were carried out with Fiji. Statistical analysis was carried out with Prism. Figures were assembled in Affinity Designer.

### Brdu/Edu labelling

For BrDU labelling experiments, BrDU (Invitrogen B23151) was injected at 50mg/kg intraperitoneally 30 min prior to harvesting embryos. For BrDU/EDU labelling experiments, BrDU was injected at 50mg/kg intraperitoneally 4 hours prior to harvesting embryos, and EdU (Thermofisher Click-iT EdU Alexa Fluor 555) was injected at 50mg/kg 30 min prior to harvesting embryos. After fixation, brain slices were incubated in 2N HCL for 30 min at 37 degrees and then washed 3 times with PBS prior to immunostaining.

### WDR81 patient cells and immunostaining

Control and WDR81 mutant primary fibroblasts were provided by Institut Imagine, Paris. The genotype of patient 1 cells was compound heterozygote 1882C>T/3713C>G and the genotype of patient 2 cells was compound heterozygote 1582C>T/4036_4041dup. Cells were grown in OPTIMEM + 10%FBS at 37 degrees in humid air containing 5% CO_2_. Fibroblast were fixed in 4% paraformaldehyde for 20 min, treated with 50mM NH4Cl for 10min, washed three times with PBS and left in a blocking solution (PBS 1% donkey serum 0.1% Triton X) for 30 min. Cells were then incubated 1 hour at room temperature with primary antibodies, washed three times in PBS and incubate for 45 min at room temperature in blocking solution with Alexa Fluor coupled secondary antibodies. Cells were then washed and mounted.

### Antibodies

Primary antibodies used: rabbit anti CUX-1 (Santa Cruz, discontinued), mouse anti Ctip-2 (Abcam ab18465), rabbit anti Pax6 (Biolegend 901301), Sheep anti TBR2/EOMES (R&D system AF6166), rabbit anti NEUN (Abcam ab177487), goat anti Phospho Histone3 (Santa Cruz SC-12927), rabbit anti BRDU (Abcam AB152095), rabbit cleaved caspase-3(CST 3661), rabbit anti Ki67 (abcam ab15580), rabbit anti EGFR (CST 4267),mouse anti p-ERK (CST 9106), rabbit anti GAPDH (Sigma AldrichG9545) and mouse anti EEA-1 (BD biosciences 610457). Secondary antibodies used: donkey Alexa Fluor 488 anti-mouse, anti-rabbit, anti-goat (Jackson laboratories 715-545-150, 711-165-152, 715-605-152), donkey Alexa Fluor 555 anti-mouse, anti-rabbit, anti-goat (Jackson laboratories 715-545-150, 711-165-152, 715-605-152), donkey Alexa Fluor 647 anti-mouse, anti-rabbit, anti-goat (Jackson laboratories 715-545-150, 711-165-152, 715-605-152).

### Expression constructs and shRNAs

For WDR81 knockdown experiments, WDR81 shRNA was provided by Genecopoeia™. The small interfering RNA sequence was ggagataagcaattggacttc and was cloned in psi-mU6.1 vector coexpressing mcherryFP. For WDR91 knockdown experiments shRNA was provided by Tebu-bio (217MSH024100-mU6). For p62 knockdown experiments shRNA was provided by Tebu-bio (217CS-MSH079315-mU6-01). Constructs were co-injected with GFP-pCagIG (Addgene 11159) at a concentration of 1ug/ul. Plasmids were introduced in the *in vivo* developing cortex by intraventricular injection and electroporation. For WDR81 rescue experiments, Cyclin D2 and Cyclin D2Thr280Ala were synthetized *in vitro* (Genescript). They were then cloned into GFP-pCagIG (Addgene 11159) after digestion by restriction enzymes EcoRI and EcoRV.

### EGF pulse assay and EGF^555^ uptake assay

For EGF pulse assay, fibroblast cultures were EGF starved for two hours before the assay. EGF was added directly to the culture medium at 0,1mg/ml. Cells were then harvested at 0, 5, 15, 30, 60, 120 minutes and processed for protein extraction. Proteins were then mixed with 4x Leammli (Biorad) and BME solution and used for Western Blot analysis. For EGF^555^ pulse assay, fibroblast cells were cultivated on glass coverslips; cells were starved for 24 hours and then exposed to at 0,1mg/ml EGF^555^ (Thermofisher E35350). Cells were fixed in paraformaldehyde 4% at 15, 30, 60, 120, 360 min and used for immunostaining.

## Acknowledgments

We acknowledge Institut Curie, member of the French National Research Infrastructure France-BioImaging (ANR10-INBS-04) and the Nikon BioImaging Center (Institut Curie, France). We thank Renata Basto, Veronique Marthiens, Iva Simeonova, Cedric Delevoye (I. Curie), Fiona Francis (IFRM), for helpful discussions and critical reading of the manuscript. J.A.C. was funded by the IC3i Institut Curie doctoral program founded by the European Union’s Horizon 2020 research and innovation program under the Marie Sklodowska-Curie actions grant agreement and from Fondation de la Recherche Medicale (FRM). A.D.B. is an Inserm researcher. This work was supported by the CNRS, I. Curie, the Ville de Paris “Emergences” program, Labex CelTisPhyBio (11-LBX-0038) and PSL.

## Author contributions

J.A.C. and A.D.B. conceived the project. J.A.C. & A.D.B. analyzed the data. J.A.C., J.B.B. & A.D.B. wrote the manuscript. J.A.C. and A.D.C. did most of the experimental procedures. J.A.C., L.D.M. and F.E.M. created the mutant mouse lines. D.A. supervised the mouse mutant colonies and performed all the crossings. L.C. and J.A.C. set up *in utero* electroporation experiments. N.B.B. provided biological samples of the affected patients A.D.B. supervised the project.

## Figure legends

**Supplemental figure 1. Genotypes and survival and WDR81**^-/-^ **mice.**

**A.** Sequencing of WDR81 knock-out animals reveals a 4 base pair deletion in KO1 and an 8 base pair deletion in KO2, both leading to frameshifts and premature STOP codons. **B**. Rate of WDR81^-/-^ embryos and pups recovered throughout time. The expected rate is 25% (dashed line). By P21, no mutant was detected.

**Supplemental figure 2. Quantification of EGFR levels and P-ERK levels for Control-2 and WDR81 patient-2 cell lines.**

**A.** Quantification of EGFR levels, normalized to control levels at T0 for Control-2 and WDR81 patient-2 cell lines (n = 5). **B**. Quantification of P-ERK levels, normalized to control levels at T5 for Control-2 and WDR81 patient-2 cell lines (n = 5). *p<0,05; **p<0,01; ***p<0,001; ****p<0,0001 by unpaired t-tests.

## Notes

### Competing Interest Statement

The authors have declared no competing interest.

