## Supplementary figures and images for "Endosomal trafficking defects alter neural progenitor proliferation and cause microcephaly"

### Supplemental figures

#### Supplemental figure 1

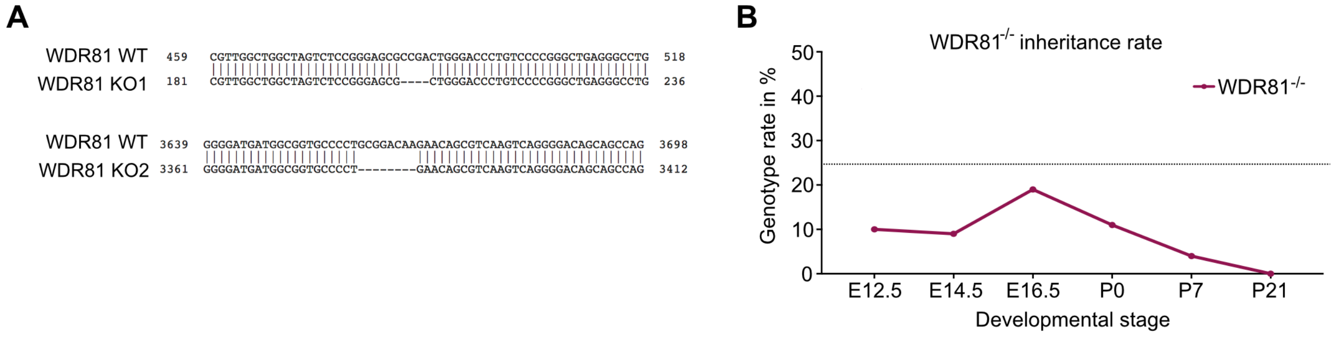

#### Supplemental figure 2

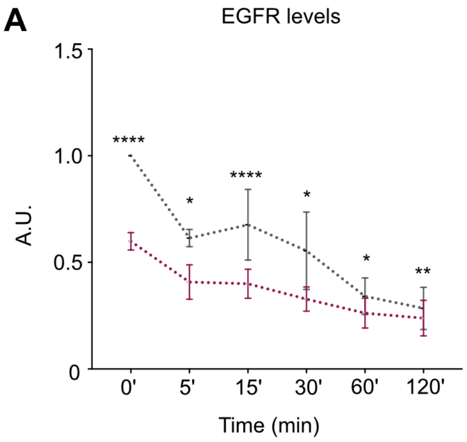


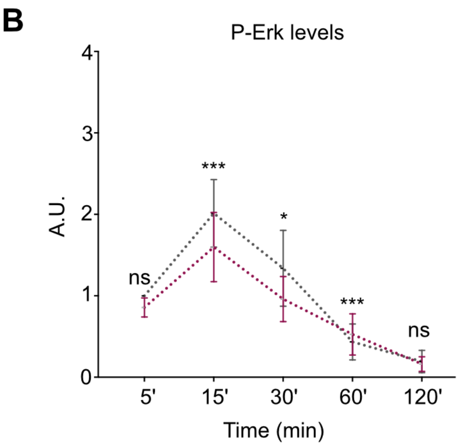
